# Attractant and repellent induce opposing changes in the chemoreceptor four-helix bundle ligand-binding domain

**DOI:** 10.1101/2023.03.29.534785

**Authors:** Lu Guo, Yun-Hao Wang, Rui Cui, Zhou Huang, Yuan Hong, Jia-Wei Qian, Bin Ni, An-Ming Xu, Cheng-Ying Jiang, Igor B. Zhulin, Shuang-Jiang Liu, De-Feng Li

## Abstract

Motile bacteria navigate toward favorable conditions and away from unfavorable environments using chemotaxis. Mechanisms of sensing attractants are well understood, however molecular aspects of how bacteria sense repellents have not been established. Here, we identified malate as a repellent recognized by the MCP2021 chemoreceptor in a bacterium *Comamonas testosteroni* and showed that it binds to the same site as an attractant citrate. Binding determinants for a repellent and an attractant had only minor differences, and a single amino acid substitution in the binding site inverted the response to malate from a repellent to an attractant. We found that malate and citrate affect the oligomerization state of the ligand-binding domain in opposing way. We also observed opposing effects of repellent and attractant binding on the orientation of an alpha helix connecting the sensory domain to the transmembrane helix. We propose a model to illustrate how positive and negative signals are generated and transduced across the membrane and built chimera proteins to illustrate a universal nature of the transmembrane signaling by the repellent.

## Introduction

Bacteria utilize flagellar motility to navigate toward or away from spatial gradients of chemical stimuli (1). This process, called chemotaxis, is vital for finding nutrients, escaping toxins, and establishing relationships with hosts (2, 3). Approximately half of known bacterial species have chemotaxis machinery genes encoded in their genomes (4), but molecular mechanism of chemotaxis is best studied in the single model organism *Escherichia coli*. where chemical signals are detected by ligand-binding domains (LBDs) of transmembrane chemoreceptors (5, 6). Chemoreceptor homodimers form mixed trimers-of-dimers (7-9), that are packed into a ternary hexagonal array with a histidine kinase CheA and a scaffolding protein CheW (10-12). The decreased concentration of an attractant chemical detected by the chemoreceptor LBD promotes CheA autophosphorylation. Phosphorylated CheA donates its phosphoryl groups to the CheY response regulator, and the phosphorylated CheY (CheY-P) interacts with the flagellar motor and triggers the clockwise rotation resulting in tumbling (1). Binding of an attractant to the chemoreceptor LBD suppresses CheA activity, which leads to dephosphorylation of CheY due to activity of its dedicated phosphatase CheZ and ultimately promotes swimming up the attractant gradient.

How small conformational changes resulting from an attractant binding propagate through the entire chemoreceptor molecule (several hundred Angströms) is not fully understood, although the role of structural and dynamic changes in various parts of the receptor was documented (13-17). In *E. coli* Tar chemoreceptor, an attractant aspartate binds at the four-helix bundle LBD dimeric interface with a stoichiometry of one molecule per LBD homodimer (18) triggering an inward sliding of the last α-helix (α4) that extends into the second transmembrane helix (TM2) (5, 19, 20). Consequently, transmembrane helices undertake piston and rotation movements and induce conformational changes in the HAMP domain that subsequently generate conformational changes in the downstream signaling domain altering CheA activity (21).

In contrast to the sensing mechanism for attractants, the molecular details of repellent recognition are poorly understood. In *E. coli*, Tar mediates a chemotactic repellent response to metal ions by an unknown mechanism (22), and no repellents that bind to Tar-LBD have been identified in large-scale screening (23). In a recent study, leucine was identified as a repellent recognized by *E. coli* chemoreceptor Tsr (24), which also senses another amino acid, serine, as an attractant. Most interestingly, leucine and serine bind to the same binding pocket and a single amino acid substitution in the binding site converts the response to leucine from repellent to attractant (24). Attractants and repellents cause the opposite behavior in chemotaxis. The *in vivo* FRET studies showed that addition of repellents increases CheA activity, whereas addition of attractants decreases the kinase activity (25, 26), implying that attractants and repellents may also trigger the opposite conformational changes in chemoreceptors. Based on the observation that the attractant binding causes the α4 helix of Tar to move towards the cytoplasm by ∼1.6 Å (27), it was proposed that repellent binding would cause an outward movement of one TM2 helix of the Tar dimer by 1-2 Å (13). While an attractant causes the chemoreceptor dimers to move apart, away from each other (26), it was suggested that a repellent might cause the dimers to move closer to each other (28). Nonetheless, these hypotheses have not been tested and how repellents trigger an opposite response from what attractants do remains unclear.

Previously, we reported that a transmembrane chemoreceptor MCP2201 in a gammaproteobacterium *Comamonas testosteroni* CNB-1 recognizes several tricarboxylic acid (TCA) cycle intermediates and mediates a positive chemotactic response towards these compounds. We also identified a binding site for an attractant citrate in the MCP2201 LBD (29), which similarly to Tar-LBD and Tsr-LBD adopts a four-helix bundle fold.

In this study, we identified malate as a repellent recognized by MCP2201. We show that malate binds to the same binding pocket that an attractant citrate, but it affects LBD dimerization and the movement of the signaling α4 helix differently. Using chimeric proteins, we further show that the signal induced by malate binding to MCP2201 LBD can be transduced to cytoplasmic signaling domains of Tar and WspA chemoreceptors, causing negative chemotaxis to malate in *E. coli* and promoting biofilm formation in *Pseudomonas aeruginosa*, respectively. These findings suggest a common transmembrane signal transduction mechanism for repellent recognition by bacterial chemoreceptors.

## Results

### Malate is a repellent recognized by MCP2201 LBD

We used several approaches to demonstrate that malate is a repellent for *C. testosteroni*, which is recognized by chemoreceptor MCP2201. In the chemical-in-plug assay (22), the chemotaxis-null mutant CNB-1Δ20, in which all chemoreceptor genes have been deleted, complemented with a plasmid carrying wild-type MCP2201 gene (CNB-1Δ20/MCP2201), swam away from agar plugs containing 10 or 20 mM L-malate a indicating that it is a repellent (Fig. 1a). In the gradient plate assay (30), where chemotactic response index (RI) values greater than 0.52 indicate an attractant and those less than 0.48 indicate a repellent (see Methods for details), CNB-1Δ20/MCP2201 strain responded to citrate with a RI value of 0.62, indicating the attractant response (confirming the previous observation (31), whereas a RI value for response to malate was 0.39, indicating the repellent response (Fig. 1b). In the transwell chemotaxis assay, which is a modified version of a classic capillary method (32, 33), more CNB-1Δ20/MCP2201 cells moved toward citrate compared to a buffer, and fewer cells moved towards increasing concentrations of malate, indicating that it acts as a repellent (Fig. 1c).

**Fig. 1.**
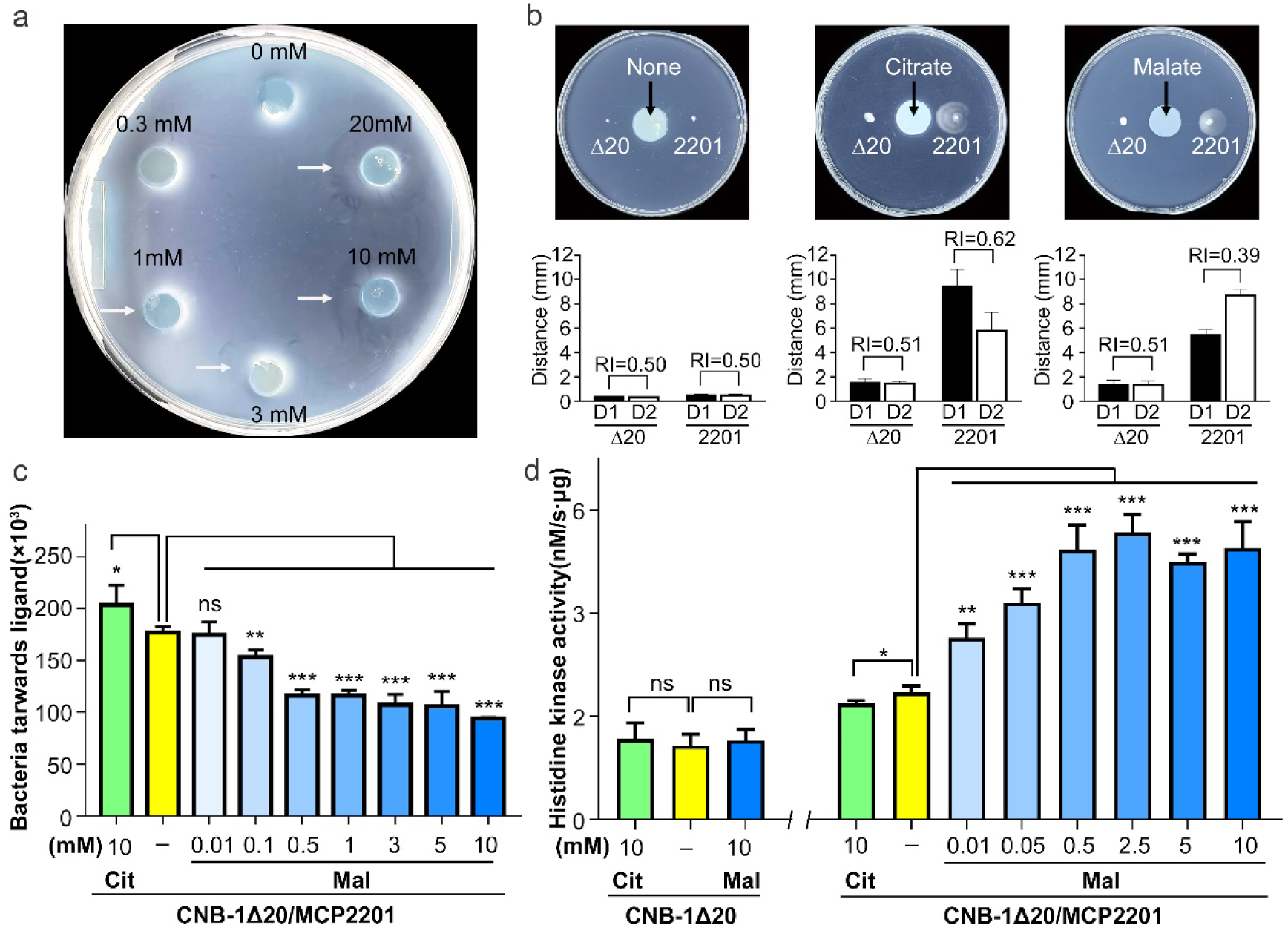
Chemotactic responses of CNB-1Δ20/MCP2201 to malate. (a) Negative chemotaxis of CNB-1Δ20/MCP2201 cells in the presence of increasing concentrations of malate measured using the chemical-in-plug method. (b) Chemotactic responses of CNB-1Δ20/MCP2201 (labeled as 2201) and CNB-1Δ20 (labeled asΔ20) to citrate (10 mM) and malate (10 mM) in soft agar plate assay. The response index (RI) was calculated as described in Materials and Methods and results are shown below each plate. Data are averages of three independent replicates. Error bars indicate standard deviations. (c) Chemotactic responses of CNB-1Δ20/MCP2201 to citrate and malate using the transwell chemotaxis assay (d) The NADH-coupled histidine kinase activity in response to citrate and malate. Results shown as the means ± SD (nL=L3). * (P<0.05), ** (P<0.01) and *** (P<0.005) showed a significantly difference between ligand-treated and non-treated groups. The “ns” stands for not significant.

Finally, we measured CheA kinase activity in response to citrate and malate. It is well established that in *E coli* attractants turn CheA activity off thereby promoting swimming, whereas repellents turn it on and promote tumbling (19, 21, 27, 34, 35). Here, the membrane fractions of CNB-1Δ20 with and without MCP2201 were extracted and subjected to NADH-coupled CheA kinase activity assay. The CheA kinase activity of strain CNB-1Δ20 was determined as 1.44 ± 0.21 nM/s·μg of membrane protein, and did not change significantly in the presence of either 10 mM citrate (1.57 ± 0.31 nM/s·μg) or 10 mM malate (1.54 ± 0.21 nM/s·μg) (Fig. 1d). The CheA kinase activity of CNB-1Δ20/MCP2201 was 2.47 ± 0.13 nM/s·μg of membrane protein, and it decreased to 2.25 ± 0.06 nM/s·μg in the presence of 10 mM citrate, but increased significantly in the presence of 0.01-10 mM of malate (Fig. 1d). Moreover, the kinase activities were fitted into a nonlinear logistic equation against L-malate concentrations. The EC50 of malate was determined as 23.32 μM, which is comparable to the binding constant of MCP2201 to malate (Kd of 18.78 μM). Thus, we concluded that malate binding to chemoreceptor MCP2201 increased CheA kinase activity as typical of repellents. Taken together, results obtained by four independent methods demonstrate that malate is a repellent, which is sensed byMCP2201 chemoreceptor.

### Repellent malate binds to the same ligand binding pocket as does attractant citrate

We previously reported that CP2201-LBD) adopts a typical four-helix bundle fold and the attractant citrate binds at an internal pocket surrounded by all four helices (29). Here, we determined the three-dimensional structure of MCP2201-LBD in complex with the repellent malate at 1.8 Å resolution and found a malate-bound MCP2201-LBD dimer in the asymmetric unit, similar to that observed in ligand-free MCP2201-LBD. As shown for the ligand-free and citrate-bound structures, the protomer of malate-bound MCP2201-LBD also folded into four helices (α1: Q59-K87, α2: A91-L118, α3: P122-A150 and α4: A154-E195). Malate was bound at an internal pocket surrounded by all four helices (Fig. 2a), which overlaps with the citrate-binding pocket (Fig. 2b). Specifically, residue R135 interacts with 1’-carboxyl group of malate via hydrogen bonds along with the potential interaction between negative and positive charges. Residue Y172 also interacts with the same carboxyl group. Residue T108 forms a hydrogen bond with the 2’-hydroxyl group of malate. Residues T105, Y138, and R142 form hydrogen bonds with the 4’-carboxyl group. Residues W71, V75 and A78 in helix α1 interact with malate via van der Waals interactions. The significant difference between malate- and citrate-binding pockets was that the ligand’s carboxyl group (coordinated by residues Y138 and R142 in both cases) forms a hydrogen bond with T105 in case of malate, but in case of citrate, this residue is not involved in ligand binding, but instead two other residues, R81 and T104, form hydrogen bonds with the ligand.

**Fig. 2.**
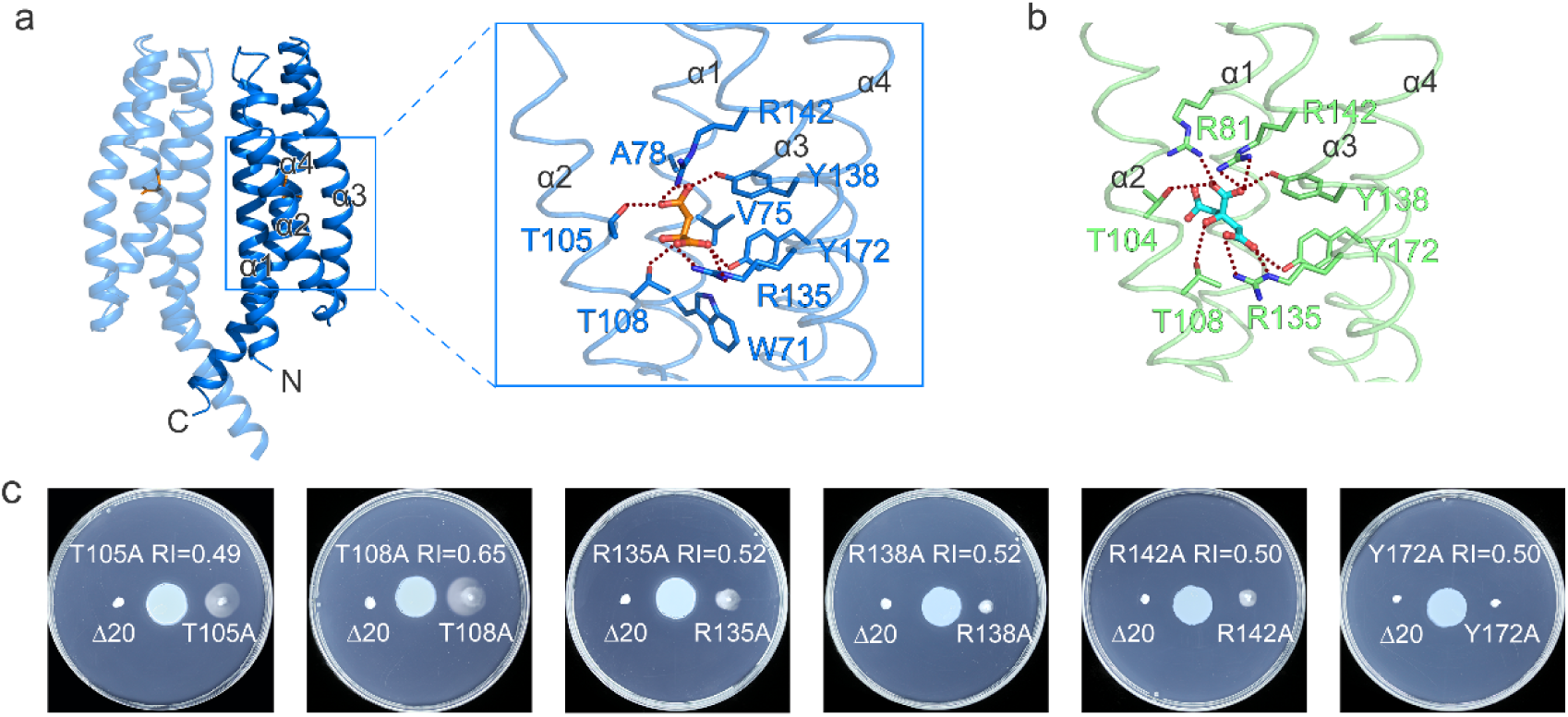
The MCP2201-LBD ligand-binding pocket and contact residues for malate and citrate binding. (a) Location of the binding pocket in MCP2201-LBD and residues involved in malate binding (brown stick representation, this study) and (b) residues involved in citrate binding (cyan stick representation, PDB accession code 6ITS). (c) Chemotactic responses to malate on the gradient soft-agar plates by CNB-1Δ20 cells harboring MCP2201 mutants. RI: T105A, 0.49; T108A, 0.65; R135A, 0.52; R138A, 0.52, R142A, 0.50, and Y172A, no response CNB-1 Δ20 used as a control).

Residues involved in malate binding were subjected to mutagenesis and the chemotaxis phenotypes of the resulted mutants were determined. The mutants T105A, R135A, R138A, R142A, and Y172A failed to respond to malate (RI range 0.49 to 0.52 (Fig. 2c)), respectively. Unexpectedly, the mutant T108A showed an attractant response to malate (moved towards malate, with RI of 0.65 Fig. 2c). We then verified malate binding affinities in T105A (no response) and T108A (inverted response) mutants and found that while the T105A mutant completely lost its ability to bind malate (the Kd was too weak to be determined), the T108A mutant bound malate with Kd of 995.1 μM (Supplementary Fig. 4). Thus, the T105A mutant showed significantly lower affinity for malate than the wild type (MCP2201-LBD (Kd of 18.78 μM) suggesting that the strength of ligand-binding to chemoreceptors might play a critical role in chemotactic responses.

### Binding of attractant and repellent differently affects the LBD oligomeric state

Previous study revealed that the ligand-free and citrate-bound recombinant MCP2201-LBDs were found as mixtures of monomers and dimers and monomers and trimers, respectively (29). In this study, we investigated the oligomeric state of MCP2201-LBDs in the presence of repellent malate. Using analytical ultracentrifugation assays, we showed that, consistently with our previous observations (29), the citrate-bound MCP2201-LBDs primarily formed monomers (major fraction, apparent molecular mass of 19 kDa), but also trimers (minor fraction, apparent molecular mass of 54 kDa) (Fig. 3a), whereas the ligand-free MCP2201-LBDs was found primarily in monomer state (large fraction, apparent molecular mass of 20 kDa), with a small fraction in dimeric (small dominant fraction, apparent molecular mass of 33 kDa) state and (Fig. 3a). In contrast, the malate-bound MCP2201-LBDs primarily formed dimers (apparent molecular mass of 34 kDa) (Fig. 3a). The oligomer disassociation constants of the ligand-free and malate-bound dimers were 32.14 μM and 0.042 μM, respectively, whereas the oligomer disassociation constant of citrate-bound trimers was hardly measureable, at approximately 10 mM (Fig. 3b). The results suggest that binding of the repellent malate and the attractant citrate promote different changes in the MCP2201-LBD oligomerization state.

**Fig. 3.**
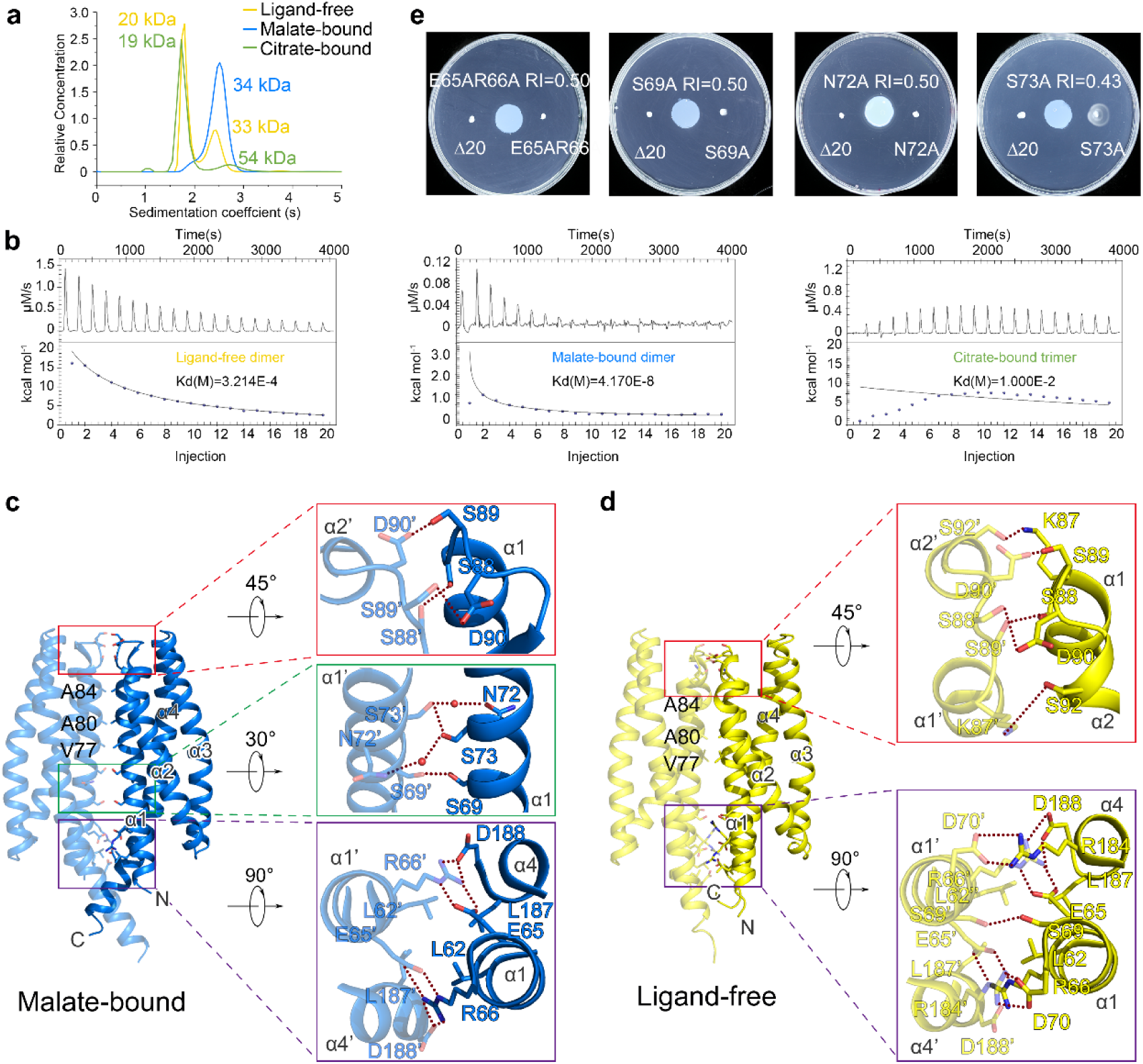
Oligomer states and dimeric interface of MCP2201-LBD. (a) Analytical ultracentrifugation assays of ligand-free, malate-bound (10 mM), and citrate-bound (10 mM) MCP2201-LBD. (b) ITC assays of MCP2201-LBD oligomer dissociation in the absence and in the presence of citrate and malate. Calorimetric dilution data (top) for injection of MCP2201-LBD (1.11 mM) without a ligand (left), in the presence of with 10 mM malate (middle), and with citrate (right) at 25°C were integrated and dilution corrected peaks were fitted to an oligomer dissociation model (bottom) to assess the dissociation constants. (c, d) The malate-bound (c, this study) and ligand-free (d, PDB accession number 5XUA) dimers and residues involved in dimeric interface. Two subunits were colored in light and dark colors. (e) Chemotactic responses to malate on the gradient soft-agar plate by CNB-1Δ20 cells harboring MCP2201 mutants.

The crystal structures of the malate-bound and ligand-free MCP2201-LBD dimers were similar, with r.m.s.d of 2.1 Å. The interface of malate-bound dimers was formed by helices α1 and α4 (Fig. 3c and 3d). Residues L62, V77, A80, A84, and L187 from two protomers contributed to hydrophobic interactions and residues E65, R66, K87, S88, S89, D90, S92, R184, and D188 contributed to hydrogen bonds at the interface (Fig. 3c). Residues S69, N72, and S73 were involved in a water-mediated hydrogen bond network at the malate-bound dimer interface, but not in the ligand-free dimer interface. The total buried area of malate-bound and ligand-free dimer was similar: approximately 1280.7 Å2 and 1281.1 Å2 per subunit, respectively. The free energies of malate-bound (-14.5 kcal/mol) and ligand-free (-10.8 kcal/mol) dimer formation calculated by PISA(36) support a more stable malate-bound dimer than the ligand-free one (Fig. 3d).

In order to evaluate the importance of the dimeric interface, we first constructed two mutants, E65A/R66A and S73A and showed that their dimer disassociation constants were significantly higher (68.86 μM for E65A/R66A and 95.48 μM for S73A (Supplementary Fig. 3)) than that of malate-bound wildtype MCP2201-LBD (0.042 μM) (Fig. 3b), indicating that the hydrogen bonds at the interface were critical for dimer formation. We then evaluated both mutants for their ability to respond chemotactically to malate, and found that the chemotactic response was diminished in S73A mutant (RI value of 0.43) and completely abolished in E65A/E66A mutant (Fig. 3e). Finally, we constructed two additional mutants, S69A and N72A, and both mutants showed no chemotactic response to malate (Fig. 3e). Taken together, these results indicated that the dimeric interface is critically important for negative chemotaxis to malate mediated by MCP2201.

### Attractant and repellent induce a swing of the LBD signaling helix in opposite directions

We superposed the ligand-free, citrate-bound, and malate-bound structures and noticed difference in positions of helices α1, α2 and the C-terminus of helix α4 (Fig. 4a and 4b). Both malate and citrate interacted with residues W71, V75, and A78 in helix α1 via Van de Waals interactions, inducing the displacement of helix α1 away from the membrane compared to the ligand-free dimer (Supplementary Fig. 1b). Malate formed hydrogen bonds with residues T105 and T108 in helix α2, whereas citrate formed hydrogen bonds with residues T104 and T108, resulting in different relative movement and bending angle variation of helices α2 (Supplementary Fig. 1a). Malate and citrate also formed hydrogen bonds with residues R135, Y138, R142, and Y172 in helices α3 and α4, however, we observed no significant conformational changes in helix α3 and in the N-terminal part of helix α4 (residues 154-182). In contrast, we observed a swing of the C-terminal part of helix α4 (residues 183-195) in ligand-bound structures compared to the ligand-free structure. Furthermore, in malate-bound and citrate-bound structures the swing was in opposing directions (Fig. 4a).

**Fig. 4.**
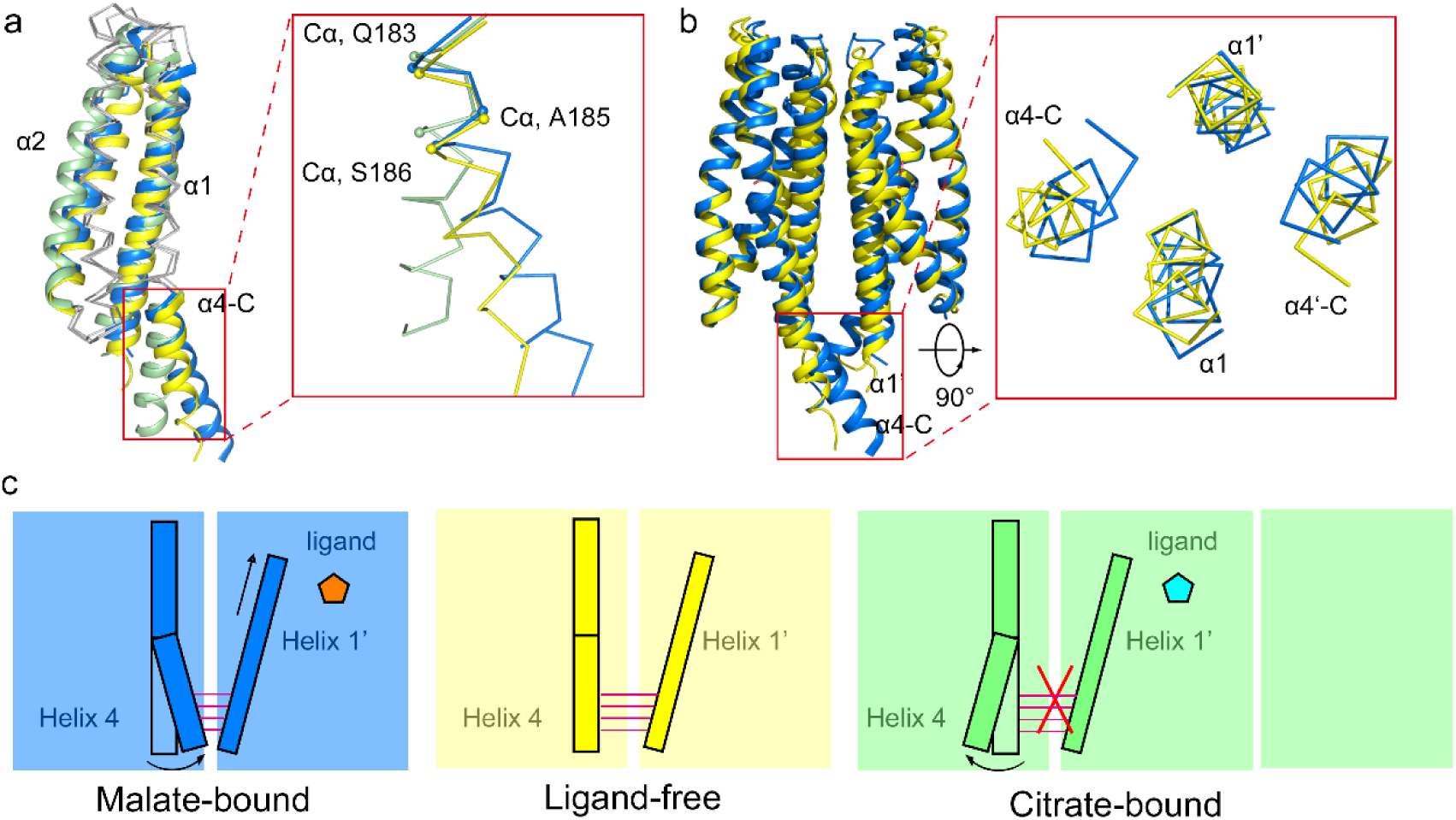
Position of C-terminus of helix. α**4 in ligand-free, malate-bound and citrate-bound MCP2201-LBD.** (a) Superposition of the ligand-free (yellow), citrate-bound (green), and malate-bound (blue) structures. The inset showed the different orientation of C-terminus of helix α4 in different structures. (b) Superposition of the ligand-free and malate-bound dimers. The inset showed the swing of C-terminus of helix α4towards helix α1 of the other subunit. (c) Chemotactic responses to malate on the gradient soft-agar plate by CNB-1Δ20 cells harboring MCP2201 mutants. (d) Cartoon representation of interaction between the C-terminus of helix α4 of one protomer and helix α1 of another protomer in ligand-free, malate-bound, and citrate-bound structures. Each color block represents a protomer.

### MCP2201 LBD enables a repellent response to malate by chimeric chemoreceptors in *E. coli* and *P. aeruginosa*

Functional chemoreceptor hybrids were successfully constructed in the past to demonstrate the common mechanism of transmembrane signaling in response to attractants by bacterial chemoreceptors and sensor histidine kinases (37). We used a similar approach to construct chimeras consisting of the MCP2201 LBD and signaling domains from well-studied chemoreceptors in model organisms - Tar, which mediates an attractant response to aspartate in *E. coli* (38), and WspA chemoreceptor which regulates biofilm formation in *P. aeruginosa* (39). The MCP2201-Tar and MCP2201-WspA chimeras were constructed by fusing the MCP2201 region containing TM1, LBD, TM2, and the HAMP domain with the signaling (kinase control) domains of Tar and WspA (Fig. 5a). *E. coli* MG1655 cells carrying the MCP2201-Tar chimera swam away from malate, with RI value of 0.37 in a gradient soft-agar swim plate assay (Fig. 5b), indicating that the transmembrane signal generated by MCP2201-LBD has been successfully transduced to the cytoplasmic domain of Tar. The MCP2201-WspA chimera was introduced into *P. aeruginosa* ΔWspA cells and the biofilm formation was evaluated by the crystal violet staining assay. Compared to the the ΔWspA strain, cells complemented with MCP2201-WspA chimera produced more biofilm in the presence increasing concentrations of L-malate (Fig. 5c), suggesting that MCP2201-WspA sensed malate and triggered the downstream biofilm formation signal.

**Fig. 5.**
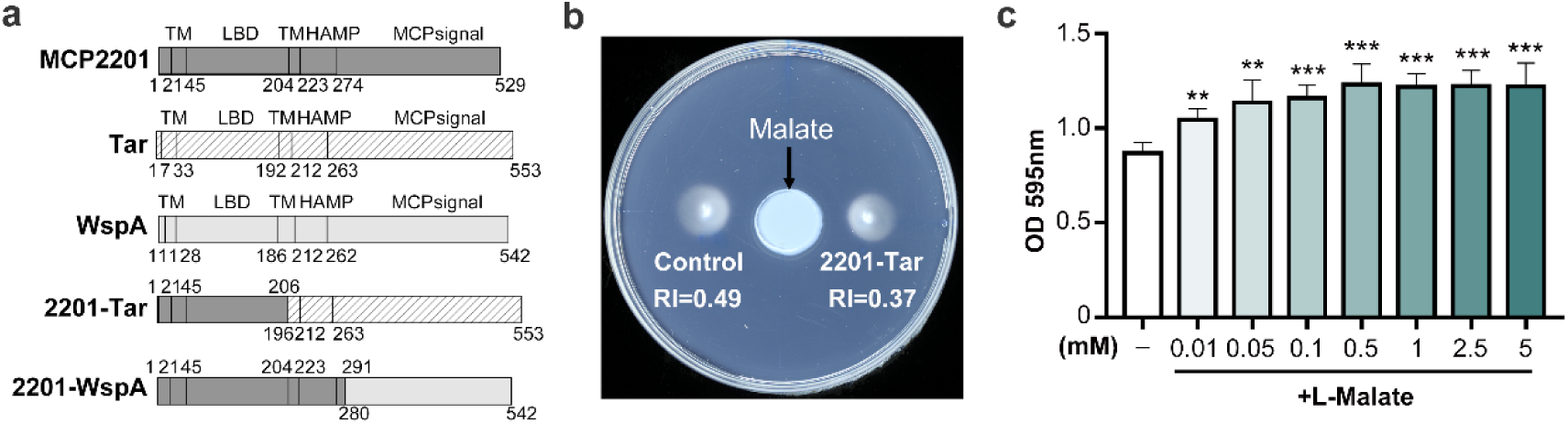
Chemotaxis of *E. coli* harboring MCP2201-Tar and biofilm formation of *P. aeruginosa* harboring MCP2201-WspA. (a) Domain arrangement of MCP2201, Tar, WspA, and MCP2201–Tar and MCP2201-WspA chimeras. (b) Chemotaxis of *E. coli* strain harboring MCP2201-Tar to malate on the gradient soft-agar plate. (c) Biofilm formation of *P. aeruginosa* strains harboring MCP2201-WspA in the presence of varying concentrations of malate assessed using crystal violet (CV) staining Results are shown as the means ± SD (nLJ=LJ3). * (P<0.05), ** (P<0.01) and *** (P<0.005) t.

**Fig. 6.**
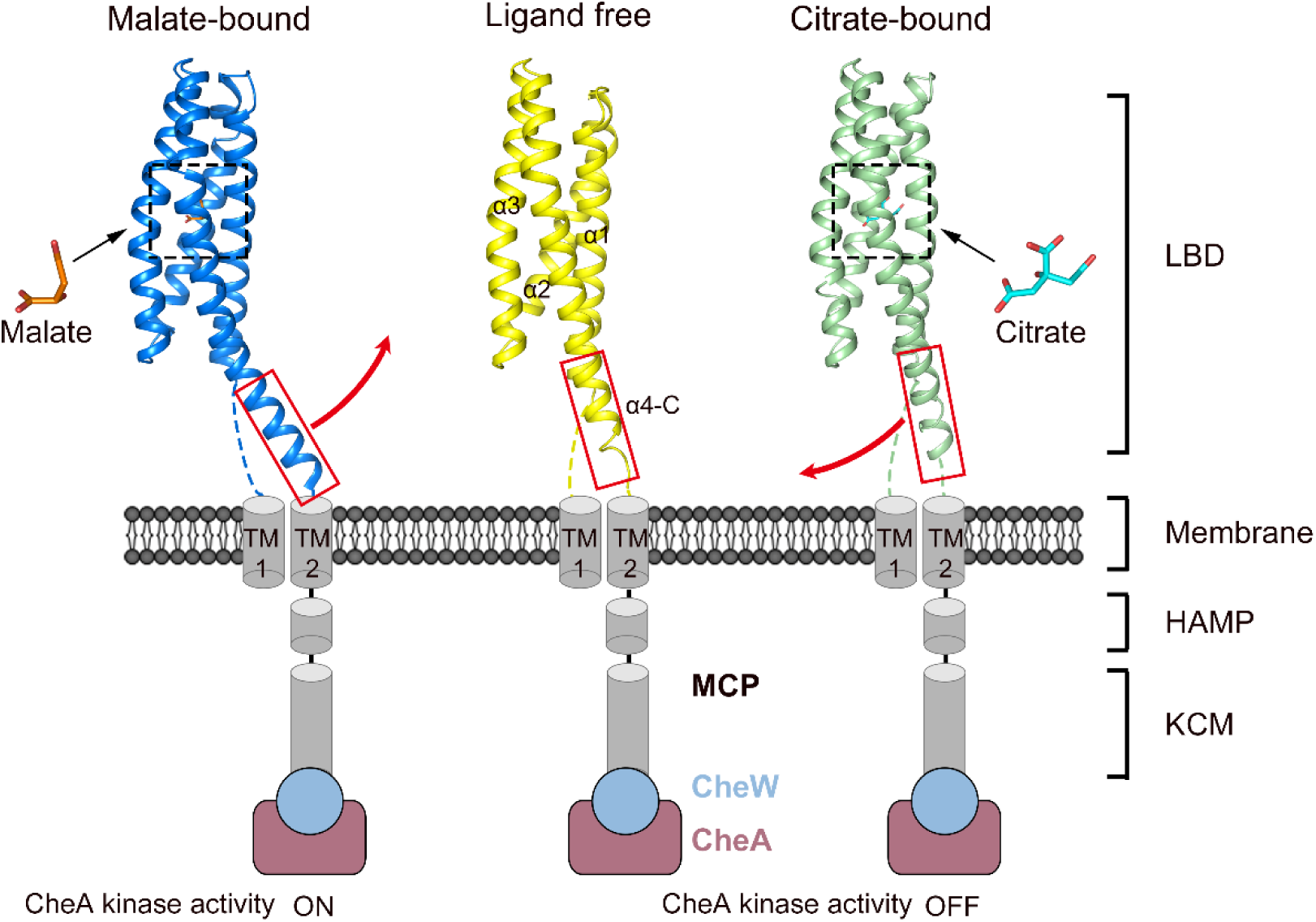
The proposed model of positive and negative chemotaxis signal generation by MCP2201. The binding of attractant (citrate) and repellent (malate) induce different orientations of helix 4 C-terminus and send opposing signals via TM2 and the HAMP domain to regulate the CheA kinase activity.

## Discussion

Molecular mechanisms of detecting chemical attractants by bacterial chemoreceptors have been studied extensively (19, 21, 27, 34). In a striking contrast, the detection mechanisms for repellents remain poorly understood. In this study, we identify malate as a repellent recognized by MCP2201 chemoreceptor of *C. testosteroni*, and demonstrate that an attractant and a repellent induce opposing changes in its four-helix bundle LBD. Finally, using chimeric chemoreceptors we show that transmembrane signaling mechanism by repellents might be universal.

We propose the following model for observed opposing conformational changes in MCP2201-LBD upon attractant and repellent binding. In the ligand-free structure, residues 183-195 in helix α4 of one protomer are packed against residues 57-70 in helix α1 of the other protomer via hydrogen bonds and Van de Waals interactions (Supplementary Fig. 1c) thus constraining their conformations and contributing to the dimeric interface. Upon malate binding, the dimer organization is not altered. Helix α1 moves away from the membrane and then induces the swing of residues 183-195 in helix α4 of the other protomer towards the direction where helix α1 moves via inter-dimer interaction (Fig. 4b). In contrast, upon citrate binding, the movement of helix α1 causes a steric hindrance that restricts dimer and favors trimer formation (Supplementary Fig. 2) (29). The citrate-bound MCP2201-LBD exists as the monomer-trimer equilibrium (29). In the monomeric and trimeric states, residues 181-195 are not involved in any interface packing thus contributing to the different orientation of the C-terminal part of helix α4 (Fig. 4c).

Several studies reported that the attractant binding alters the oligomeric state of chemoreceptor LBDs, including the 4HB_MCP domains of *E. coli* Tar (18, 40-42), *Pseudomonas putida* PcaY_(43) and *P. aeruginosa* CtpH (44), the HBM domains of *P. putida* McpS (45) and McpQ (46), and the PilJ domain of *P. aeruginosa* McpN (47), all of which adopt the four-helix bundle fold. It was suggested that ligand binding at the dimeric interface alters the oligomeric state because ligands interact with both subunits. On the other hand, no such alteration was seen in Cache domains (48) where attractants bind to a defined pocket in each subunit. For example, in *P. aeruginosa* the dCache_1 domain of PctA is found as a monomer in both ligand-free and ligand-bound states (49) and the sCache_2 domain of PA2652 exists as a dimer in both states (50). MCP2201-LBD represents the third kind of chemoreceptor binding domains, where ligands do not bind at the dimeric interface of the four-helix bundle LBD, but do alter its oligomerization. We found the attractant binding shifts the ligand-free weak MCP2201-LBD dimer state (an equilibrium of monomer and homodimer states) to the weak trimer state (an equilibrium of monomer and homotrimer states), whereas the repellent binding shifts it to the stable dimer state.

The major structural difference between ligand-free, citrate-bound, and malate-bound MCP2201-LBD include a piston movement of helix α1, a bending of helix α2, and a swing movement of C-terminus of helix α4. We propose that the malate binding induces the displacement of helices α1, modifies the dimeric interface, increases the dimer formation and then alters the orientation of the helix α4 C-terminus (via the dimeric interface interaction between helix α1 in one protomer and α4 in the other protomer). In contrast, the citrate binding disrupts the dimeric interface, thereby increasing the conformational freedom of residues involved in ligand-free and malate-bound dimer formation and altering the orientation of the C-terminus of helix α4.

Differences in ligand-induced conformational changes reported in *E. coli* Tar and Tsr LBDs and observed here in MCP2201-LBD are not necessarily surprising. All three domains belong to the same superfamily, which the leading protein domain database InterPro (51) classifies as “Four helix bundle sensory module for signal transduction” (InterPro accession cl0457). However, within this superfamily, MCP2201 LBD belongs to the largest family that represents ∼50K of proteins and ∼14K of bacterial and archaeal species (InterPro accession IPR024478), whereas Tar and Tsr LBDs are found in a smaller family of ∼15K of proteins and ∼4K of species (InterPro accession IPR003122). InterPro protein families are defined by sequence similarity, thus MCP2201-LBD and Tar/Tsr LBDs are only distantly related despite sharing the same fold.

Despite remote homology, Tsr and MCP2201 share a remarkable common property. Using molecular docking and mutation experiments, a recent study revealed that the repellent leucine binds to the same binding site on Tsr-LBD as the attractant serine, and only minor changes in one or two amino acid residues in the LBD determine whether the ligand induces attractant or repellent response (24). The single amino acid substitution in the binding pocket reversed the response to leucine from negative to positive chemotaxis (24). Here, we reveal a similar case, where intermediates of the TCA cycle citrate and malate, bind to the same binding site of MCP2201-LBD causing an attractant and a repellent response, respectively. Furthermore, as in the case of Tsr response to leucine, a single amino acid substitution in the binding pocket of MCP2201-LBD converted the response to malate from negative to positive chemotaxis. This apparent similarity is striking, because the location of binding sites is quite different: serine and leucine bind at the dimer interface of Tsr-LBD, whereas citrate and malate bind inside the MCP2201-LBD monomer.

Over a hundred of different LBD types were identified in bacterial chemoreceptors (52), but all chemoreceptors contain only one type of a signaling domain (53), and membrane topology Class I, where extracytoplasmic LBD is flanked by two TM helices, is predominant in chemoreceptors (54) and widespread in sensor histidine kinases. Consequently, signal transduction from extracytoplasmic LBDs to signaling domains involves their shared structural elements – a helix adjacent to TM2, TM2 itself, and the downstream HAMP domain (13). Taking advantage of this common transmembrane signaling arrangement, various chimera proteins containing LBDs of different types fused with heterologous chemoreceptor signaling domains were successfully constructed and shown to confer novel sensory functions on various bacterial species. For example, a fusion of NarX-LBD, which binds nitrate, and PctC-LBD, which binds GABA, with Tar signaling domain enabled positive chemotaxis in *E. coli* to nitrate and GABA, respectively (37, 55). Functional heterologous chimeras of the *P. aeruginosa* WspA chemoreceptor, which controls biofilm formation were also constructed by fusing its signaling domain with LBDs from amino acid sensing chemoreceptors (39).

In this study, we constructed the MCP2201-Tar and MCP2201-WspA chimeras containing the ligand-binding domain of MCP2201 and the signaling domains of Tar or WspA. These chimeras successfully conferred the ability to perform negative chemotaxis to L-malate on *E. coli* cells and the ability to upregulate the biofilm formation in the presence of L-malate on *P. aeruginosa* cells, suggesting that responses initiated by repellents share the common transmembrane signaling mechanism.

## Materials and Methods

### Bacterial strains, plasmids, media and growth conditions

Bacterial strains and plasmids used in this study are listed in Table S2. Genetic complementation in *Comamonas testosteroni* CNB-1 and *Escherichia coli* MG1655 was conducted using pBBR1MCS-2 and that in *Pseudomonas aeruginosa* ΔWspA using pHERD20T. *Comamonas testosteroni* CNB-1 was cultivated in LB broth at 30°C, and *Escherichia coli* MG1655 and *Pseudomonas aeruginosa* ΔWspA strains in LB broth at 37°C.

### Site-directed mutagenesis and chimera construction

The DpnI mediated site-directed mutagenesis was conducted as previously described (29). The MCP2201-Tar and MCP2201-WspA chimeras were constructed using overlapping PCR method, similar to that used for NarX-Tar chimera (37).

### Chemical-in-plug assay

Gradient soft-agar swim plate assay were performed as previously described with minor changes (22). Briefly, chemoeffector-containing agar plugs were prepared by mixing molten 1.5% (wt/vol) agar with 0, 0.3, 1, 3, 10, 20 mM malate and placed into the Petri dish. *Comamonas testosteroni* CNB-1 was grown in LB medium to OD_600_ of 0.5-0.7, washed and re-suspended in Chemotaxis buffer (40mM NaH2PO4, 10uM EDTA, 0.05% glycerol, pH 7.5) and MSB medium (1 g/L Na_2_HPO_4_·12H_2_O, 0.5 g/L KH_2_PO_4_, 0.03 g/L MgSO_4_·7H_2_O and 1 g/L NH_4_CL) respectively. Bacterial suspensions were poured into Petri dishes andincubated at room temperature for 2–10 min and then examined to identify the clearing zone around the plug.

### Gradient soft-agar swim plate assay

Semisolid-agar assay were performed as previously described(30, 56). Briefly, chemoeffector-containing agar plugs were prepared by mixing molten 1.5% (wt/vol) agar with 10 mM L-malate or citrate and placed into the center of the Petri dish containing the semisolid-agar with 0.25% (wt/vol) agarose in MSB medium (1 g/L Na_2_HPO_4_·12H_2_O, 0.5 g/L KH_2_PO_4_, 0.03 g/L MgSO_4_·7H_2_O and 1 g/L NH_4_CL). *C. testosteroni* CNB-1 cells were grown in LB medium to OD_600_ of 0.8, washed and re-suspended in 50 μl MSB medium. 0.5 μl of bacterial suspensions were inoculated 2 cm away from the center of the ligand plug. The assay plates were incubated at 30°C for 24 h. For *E. coli* MG1655 cultures, the media for plate assay contained 10 mM KPO_4_ (pH 7.0), 1 mM (NH_4_)_2_SO_4_, 1 mM MgCl_2_, 1 mg/L thiamine HCl, 0.1 mM threonine, methionine, leucine, and histidine andcells were inoculated 2.5 cm away from the center of the plug and incubated at 37°C for 24 h. The distances from the inoculation sites to the colony edges closest to (D1) and furthest from (D2) the agar plug center were measured and the response index (RI) values were calculated as follows: RI = D1/(D1 + D2), as described previously (30). Attractant and repellent were defined when RI values greater than 0.52 and less than 0.48, respectively.

### Transwell chemotaxis assay

Transwell chemotaxis assay were performed as previously described (32, 33). The experimental setup consists of a cylindrical top well insert with transparent PET membrane placed in 24-well plate. Briefly, *C. testosteroni* CNB-1 cells were grown in LB medium to OD600 of 0.5-0.7, washed and re-suspended in chemotaxis buffer (40mM NaH2PO4, 10uM EDTA, 0.05% glycerol, pH 7.5) or MSB medium (1 g/L Na2HPO4·12H2O, 0.5 g/L KH2PO4, 0.03 g/L MgSO4·7H2O and 1 g/L NH4CL) respectively. 700 µl of MSB medium containing different concentrations of citrate or malate were added to 24 well cell culture plates. Add 300 µl of the bacterial suspensions into the top well. After incubation at 30 °C for 60 min, the number of cells that entered each well was calculated (in CFUs/mL) by taking into account the dilution factor.

### Protein expression and purification

The coding sequence for MCP2201-LBD (residues 58-203) was cloned in pET22b expression vector (Novagen) with an N-terminal His_6_-tag. The constructed plasmids were transformed into *E. coli* BL21 (DE3). The cells were cultured in LB medium with 50 μg/ml kanamycin at 37°C to an OD_600_ of 0.8-1.0. Protein expression was induced with 0.2 mM IPTG at 25°C for 10 h. Bacterial cells were harvested by centrifugation at 5,000 g for 30 min. The cell pellet was re-suspended in lysis buffer consisting of 50 mM Tris buffer, pH 7.5, 200 mM NaCl and 10 mM imidazole. The cells were lysed on ice by sonication and then centrifuged at 20,000 g for 30 min. The supernatant was incubated with nickel affinity resins (Ni-NTA, Qiagen) at 4°C for 30 min. These resins were washed three times with washing buffer containing 20 mM Tris pH 7.5, 1 M NaCl and 20 mM imidazole. Protein was eluted with elution buffer containing 20 mM Tris pH 7.5, 200 mM NaCl, and 250 mM imidazole. The eluted protein was loaded on a HiTrap Q HP column (GE Healthcare) in buffer consisting of 20 mM Tris, pH 7.5 and 150 mM (SEC buffer) and eluted using SEC buffer supplied with 1 M NaCl. The elution was then loaded on a Superdex 200 10/300 GL column (GE Healthcare) and eluted in the SEC buffer. The fractions containing pure protein were concentrated to 10 mg/ml for crystallization screening in SEC buffer.

### Crystallization, data collection and structure Determination

L-malate-bound MCP2201LBD crystals were obtained using the hanging-drop vapor diffusion method at 289 K, by mixing equal volumes of protein (10 mg/ml) and reservoir solution that contained 0.2 M Ammonium acetate, 0.1 M Bis-Tris pH 5.6, 22% PEG 3,350, 5%Glycerol. The best crystals were transferred to mother liquor containing 20% glycerol as a cryoprotectant and then flash-frozen in liquid nitrogen. The diffraction datasets were collected at beamline BL19U1, Shanghai Synchrotron Radiation Facility (SSRF, China). The data were processed and scaled with the iMOSFLM(57), XDS(58) and/or CCP4 suite(59). The structure was solved by molecular replacement with Phaser in the PHENIX suite(60) and the structure of ligand-free MCP2201LBD (PDB entry 5XUA) was employed as a search model. The L-malate-bound MCP2201LBD structure was built by PHENIX AutoBuild and refined with PHENIX and Coot(61). The data collection and refinement statistics are summarized in Table S1. Figures were prepared with PyMOL (http://www.pymol.org).

### Analytical ultracentrifugation assay

MCP2201-LBD was diluted to 75 μM in SEC buffer with or without 10 mM ligand.Sedimentation velocity analytical ultracentrifugation experiments were performed on a Beckman XL-I analytical ultracentrifuge (Beckman Coulter, Brea, CA, USA) at 4 °C with a rotor speed of 60,000 rpm for 7 h. Results were analyzed using Sednterp and the sedimentation coefficients and apparent molecular masses were calculated as previously described (62).

### Isothermal titration calorimetry

ITC experiments were performed at 25 °C using Affinity ITC (TA Instruments). For dimer dissociation assays, 1.1 mM proteins dissolved in SEC buffer with or without 10 mM ligand, were injected into the sample cell containing the identical buffer mixture. For ligand-binding assays, 100 μM proteins were introduced into the sample cell and titrated with ligand dissolved in SEC buffer. Data were analyzed with the NanoAnalyze software package using independent model (TA Instruments, New Castle, USA).

### NADH-coupled ATPase assay

The ATP hydrolysis abilities of membrane fractions were used to access the histidine kinase activities in the presence and in the absence of MCP2201. The NADH-coupled ATPase assay was performed as previously described (63) with some modifications. The membrane fraction was added to 200 μl reaction mixture contained 85 mM HEPES-KOH pH 8.0, 85 mM KCl, 6.5 mM MgAC_2_, 1 mM phosphoenolpyruvate, 1 mM ATP, 2 mM NADH, 23 units of pyruvate kinase and 10 units of lactate dehydrogenase with or without ligand. Abs 340 was determined using UV2802 spectrophotometer (UNICO). The ATP hydrolysis rates were calculated with the NADH molar absorption coefficient of 6220 L mol^-1^ cm^-1^. Experiments were repeated at least three times.

### Biofilm formation assay

Biofilm formation assay was performed following a previously published protocol with minor modifications (64). Strain *P. aeruginosa* ΔWspA mutant complemented with MCP2201-WspA chimera strain was grown overnight in LB, washed and diluted to the OD 600 of 0.008 in Jensen medium. The effect of L-malate on biofilm formation was examined by incubation with varying concentrations of L-malate (0, 10μM, 50μM, 100μM, 500μM, 1mM, 2.5mM, 5mM) in the diluted cells. 100 μl aliquots of the diluted cells were pipetted into wells of a sterile 96-well U-bottom microtiter plate. After incubation at 30 °C for 24 h without shaking, attached cells were washed with ddH_2_O, and then stained with 0.1% crystal violet solution. The crystal violet was then dissolved in 200 μl of 40% acetic acid and its absorbance at 595 nm was measured. Experiments were repeated for three times.

## Acknowledgements

We are grateful to Prof. Lvyan Ma at the Institute of Microbiology, Chinese Academy of Sciences for kindly providing strain *Pseudomonas aeruginosa* ΔWspA and plasmid pHERD20T. We thank the staff members at the BL19U1 beamline at the National Center for Protein Sciences Shanghai and the Shanghai Synchrotron Radiation Facility, Shanghai, China, for technical assistance with the data collection. We also thank Dr. Wei Zhang of the Institute of Microbiology, CAS, for assisting with the ITC experiments. This work was supported by grants from the National Natural Science Foundation of China (92051101 and 31870037 to D.-F.L.), National Key R&D Program of China (2019YFA0905500 to S.-J. L and D.-F.L.), the program Youth Innovation Promotion Association CAS (2014079 to D.-F.L.) and the US National Institutes of Health (R35GM131760 to I.B.Z.).

## Author contributions

S.-J.L. and D.-F.L. conceived the study and designed the experiments; D.-F.L., L.G. and Y.H. performed the crystal structure determination; Y.-H.W., R.C., Z.H., L.G., J.-W.Q., B.N., A.-M.X., and C.-Y.J. performed the mutations, biochemical experiments and phenotypic assays; and S.-J.L., L.G., I.B.Z. and D.-F.L. analyzed the data and wrote the manuscript. All authors discussed the results and commented on the manuscript.

## Competing interests

All authors declare no competing interests.

## Data availability

The structure factor and coordinate files have been deposited in the Protein Data Bank under the accession codes number 7WRM.

## Notes

### Competing Interest Statement

The authors have declared no competing interest.

