## Supplementary Figures and Tables for "Attractant and repellent induce opposing changes in the chemoreceptor four-helix bundle ligand-binding domain"

| **Contents** | **Page** |
| --- | --- |
| Supplementary Table 1 | 2 |
| Supplementary Table 2 | 3 |
| Supplementary Figure 1 | 4 |
| Supplementary Figure 2 | 5 |
| Supplementary Figure 3 | 6 |
| Supplementary Figure 4 | 7 |
| Supplementary Figure 5 | 8 |

**Supplementary Table 1 Data collection and refinement statistics**

|  | MCP2201LBD (L-malate-bound) |
| --- | --- |
| **Data collection** |  |
| Space group | P 2_1_ 2_1_ 2_1_ |
| Cell dimensions  a, b, c (Å)  α, β, γ (°) | 47.42, 59.84, 98.06  90.00, 90.00, 90.00 |
| Resolution (Å) | 42.70–1.80 (1.90–1.80) |
| R_merge_ | 0.101 (0.698) |
| I/σ(I) | 16.5 (3.5) |
| CC_1/2_ | 99.8 (88.5) |
| Completeness (%) | 98.8 (97.6) |
| Redundancy | 11.8 (10.7) |
| **Refinement** |  |
| No. reflections | 26185 |
| R_work_/R_free_ | 0.1739 / 0.2053 |
| No. Non-H atoms |  |
| Protein | 2187 |
| Ligand/ion | 26 |
| Water | 217 |
| B factors |  |
| Protein | 26.1 |
| Ligand/ion | 22.5 |
| Water | 37.7 |
| R.m.s. deviations |  |
| Bond lengths (Å) | 0.006 |
| Bond angles (°) | 0.804 |
| Ramachandran plot |  |
| Favored | 97.8% |
| Allowed | 2.2% |
| Outliers | 0.0% |

Values in parentheses refer to the highest resolution shell.

**Supplementary Table 2 Strains and plasmids used in this study**

| Strains/plasmids | Relevant genotype or description | Source |
| --- | --- | --- |
| **Strains** |  |  |
| *Comamonas testosteroni* |  |  |
| CNB-1 |  | ^1^ |
| CNB-1Δ20 | All putative chemoreceptor genes were disrupted in strain CNB-1 | ^2^ |
| *Escherichia coli* |  |  |
| DH5α | F^-^ *φ*80d *lacZ*ΔM15 Δ (*lacZYA-argF*) U169 *rec*A1 *end*A1 *hsd*R17(r_K_^-^ m_K_^+^) *sup*E44 λ- *thi*-1 *gyr*A96 *rel*A1 *pho*A; host for DNA manipulations | ^3^ |
| BL21(DE3) | F^-^ *omp*T *hsd*S_B_(r_B_^-^ m_B_^-^) *gal* *dcm* (DE3) | Novagen |
| *P. aeruginosa*  PAO1  ΔWspA | Wild-type strain  *wspA* in-framedeletion in PAO1 | ^4^  ^4^ |
| **Plasmids** |  |  |
| pBBR1MCS-2 | Km^r^, *lacPOZ*’ broad host vector with R type conjugative origin | ^5^ |
| pBBR1MCS2-MCP2201-E65AR66A | Carries MCP2201 with E65AR66A mutation | This work |
| pBBR1MCS2-MCP2201-S69A | Carries MCP2201 with S69A mutation | This work |
| pBBR1MCS2-MCP2201-N72A | Carries MCP2201 with N72A mutation | This work |
| pBBR1MCS2-MCP2201-S73A | Carries MCP2201 with S73A mutation | This work |
| pBBR1MCS2-MCP2201-S73F | Carries MCP2201 with S73F mutation | This work |
| pBBR1MCS2-MCP2201-S73R | Carries MCP2201 with S73R mutation | This work |
| pBBR1MCS2-MCP2201-V77W | Carries MCP2201 with V77W mutation | This work |
| pBBR1MCS2-MCP2201-A80F | Carries MCP2201 with A80F mutation | This work |
| pBBR1MCS2-MCP2201-A80W | Carries MCP2201 with A80W mutation | This work |
| pBBR1MCS2-MCP2201-A84F | Carries MCP2201 with A84F mutation | This work |
| pBBR1MCS2-MCP2201-A84W | Carries MCP2201 with A84W mutation | This work |
| pBBR1MCS2-MCP2201-Tar | Carries MCP2201 spanning residues 1-206 and Tar spanning residues 196-553 | This work |
| pHERD20T |  |  |
| pHERD20T- MCP2201-WspA | Carries MCP2201 spanning residues 1-291 and WspA spanning residues 280-542 | This work |
| pET22b |  |  |
| pET22b-*mcp2201*LBD | pET22b derivative for expression of MCP2201LBD, spanning residues 58-203 and used for L-malate-bound form structure determination. | This work |

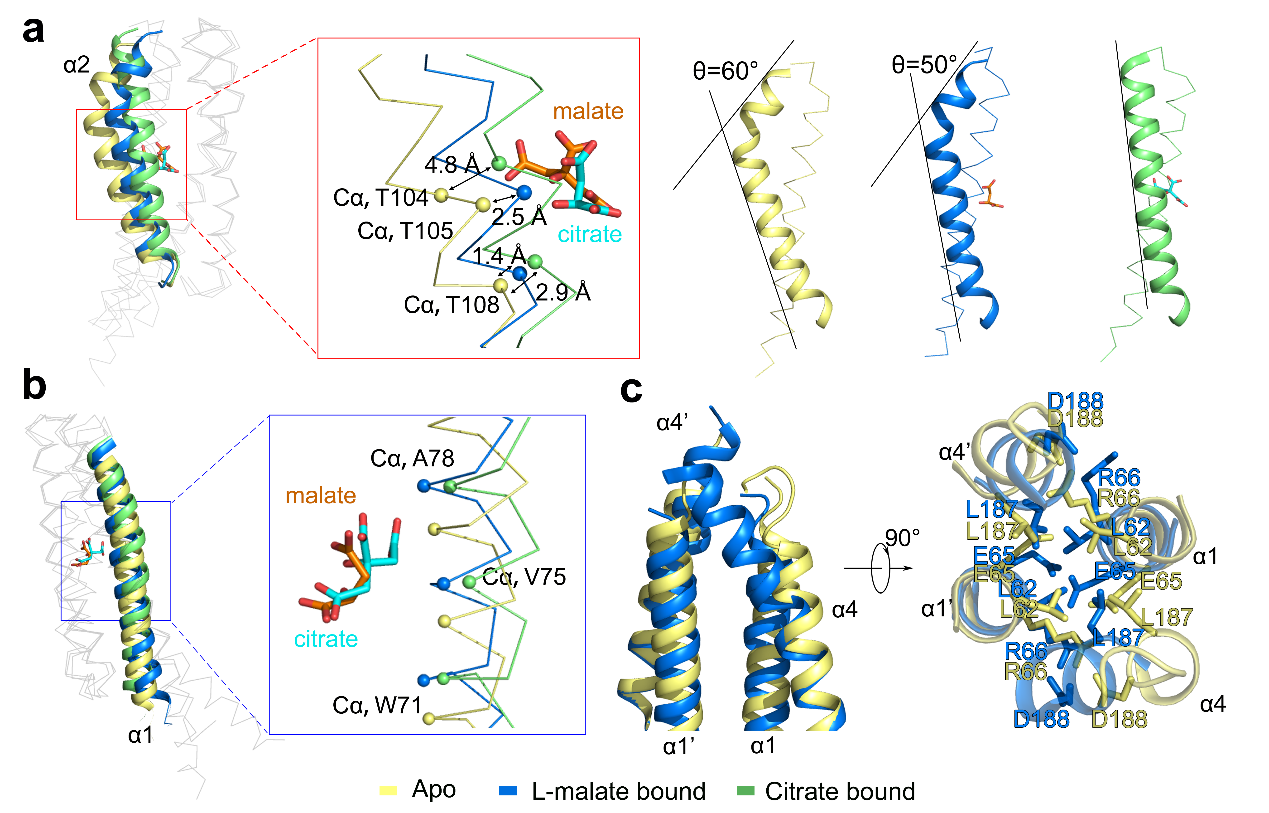

**Supplementary Figure 1. Structural comparison of apo-, citrate-bound and L-malate-bound MCP2201-LBD.**

(a) The movements of residues T104, T105, and T108 of helix 2 towards the ligand and the bend angles of the C-terminus of helix α2 in apo- (yellow) , L-Malate-bound (blue) and Citrate-bound (green) structures. (b) The displacements of helix 1 away from the membrane upon ligand binding. (c) The packing of the N-terminus of helix 1 and the C-terminus of helix 4 in the dimeric interface.

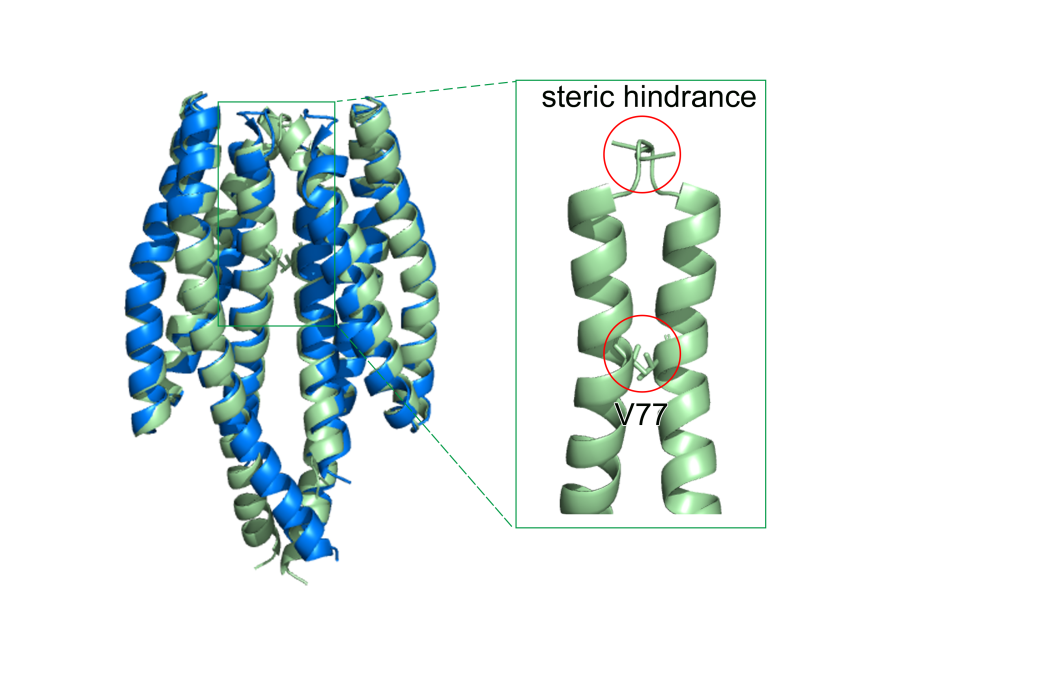

**Supplementary Figure 2. The binding of citrate induces steric hindrance that prevents the dimerization of MCP2201LBD.** Two citrate-bound MCP2201LBD monomer (green) is superposed with a L-Malate-bound dimer (blue). There are two steric hindrances (red cycles), one is Val77 and the other one is the loop between helix α1 and α2.

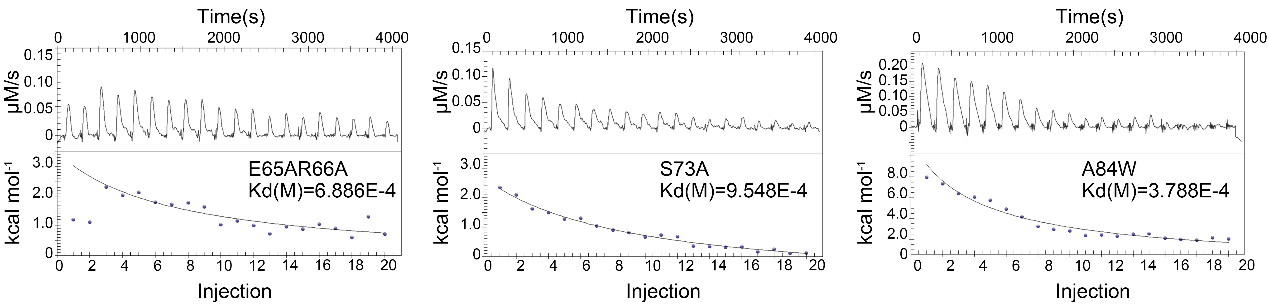

**Supplementary Figure 3. The effect of mutations in the dimeric interface on dimer dissociation.**

Calorimetric dilution data for the dissociation of E65AR66A, S69A, A84W mutant MCP2201LBD dimers in the presence of L-malate. Raw titration data for injection of MCP2201LBD (1.11 mM) into 10 mM L-malate solution at 25 °C are listed at the top, integrated and dilution corrected peaks are fit to a dimer dissociation model and listed at the bottom.

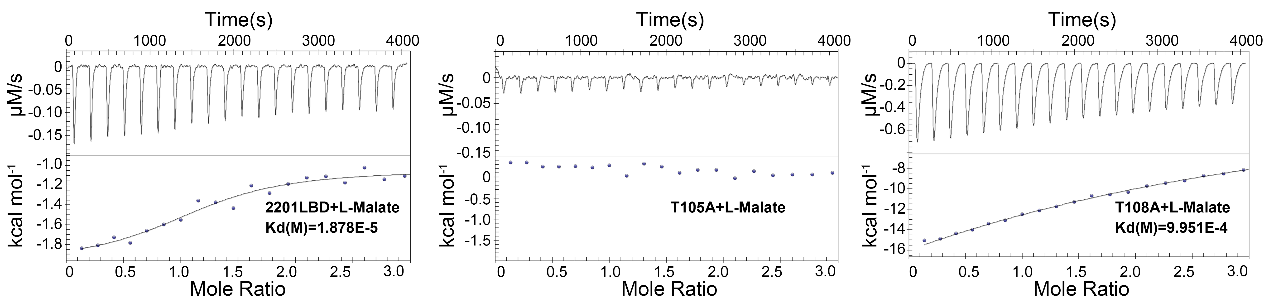

**Supplementary Figure 4. The effect of mutations in the binding pocket on L-malate binding.**

Typical raw titration curve of MCP2201LBD and T105A, T108A mutant (0.1 mM) for L-malate (1 mM) binding at 25 °C are listed at the top, integrated and normalized heat signals versus molar ratio are listed at the bottom.

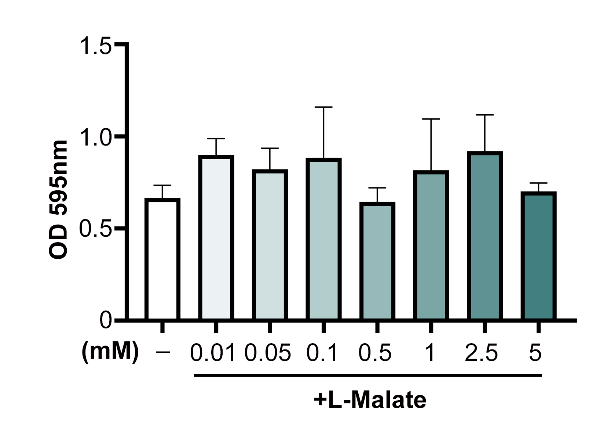

**Supplementary Figure 5. The biofilm formation ability of *P. aeruginosa.*** Bioﬁlm formation of *P. aeruginosa* strains harboring wild type WspA assessed by crystal violet (CV) staining supplemented with different concentrations of malate. Data are shown as the means ± SD (n = 3).
